# Drought duration determines the recovery dynamics of rice root microbiomes

**DOI:** 10.1101/2020.09.29.314781

**Authors:** Christian Santos-Medellin, Zachary Liechty, Joseph Edwards, Bao Nguyen, Bihua Huang, Bart C. Weimer, Venkatesan Sundaresan

**Author notes:** Department of Plant Pathology, University of California, Davis, Davis, California, USA. Department of Integrative Biology, University of Texas, Austin, Texas, USA. Microbiology and Environmental Toxicology Department, University of California, Santa Cruz, Santa Cruz, California, USA. Contributed equally to this work (Christian Santos-Medellin, Zachary Liechty).

## Abstract

As extreme droughts become more frequent, dissecting the responses of root-associated microbiomes to drying-wetting events is essential to understand their influence on plant performance. Here, we show that rhizosphere and endosphere communities associated with drought-stressed rice plants display compartment-specific recovery trends. Rhizosphere microorganisms were mostly affected during the stress period, whereas endosphere microorganisms remained altered even after irrigation was resumed. The duration of drought stress determined the stability of these changes, with more prolonged droughts leading to decreased microbiome resilience. Drought stress was also linked to a permanent delay in the temporal development of root microbiomes, mainly driven by a disruption of late colonization dynamics. Furthermore, a root-growth-promoting *Streptomyces* became the most abundant community member in the endosphere during drought and early recovery. Collectively, these results reveal that severe drought results in enduring impacts on root-associated microbiomes that could potentially reshape the recovery response of rice plants.

## Background

Drought is the largest contributor to world-wide crop loss [1, 2]. In the United States alone, major drought events between 2011 and 2018 resulted in agricultural losses totaling 78 billion dollars [3]. With an average of 25% yield reduction under drought [4], rice is particularly susceptible to this abiotic stress, due in part to its semi-aquatic growth habit and its small root system [5]. Rice responds to drought episodes through a slew of molecular, physiological, and morphological changes aimed to mitigate stress and facilitate recovery after rewatering [6]. Plant-microbe symbioses can further boost stress resistance by enhancing the plant response to environmental perturbations [7]. As such, harnessing plant-microbe interactions has emerged as a complementary approach to reduce crop losses associated with drought [8] and understanding the ecological principles governing root microbiome assembly under environmental stressors has become a research priority [9–11].

Drought triggers a compartment-specific restructuring of the rice root microbiota, with endosphere communities displaying a more pronounced response than rhizosphere communities [12]. This compositional shift is characterized by a prominent increase of a diverse group of monoderm bacteria, including Actinobacteria, Chloroflexi, and aerobic Firmicutes. Such taxonomic signatures are consistent across multiple rice cultivars and soil types. Similar trends have been independently observed in a wide variety of plant species, across cereals and dicots [13, 14], indicating that monoderm enrichment is a phylogenetically conserved response in plants under drought stress. While these cross-sectional studies have shed light on the compositional changes that root-associated microbiomes undergo during drought, the temporal dynamics upon rewatering are less understood. This recovery period is particularly relevant as both plants and microbes undergo quick physiological changes that can reshape the underlying network of biotic interactions [8].

Since rhizosphere and endosphere communities undergo compositional changes throughout the life cycles of their hosts [15–20], the temporal development of root microbiomes should also be considered when investigating community dynamics in response to drought [17, 18]. In irrigated rice, rhizosphere and endosphere communities display a highly conserved temporal development characterized by a rapid turnover during the early vegetative stages followed by a relative stabilization as the host transitions into flowering [17, 20]. These community dynamics are driven by a phylogenetically diverse group of microbial taxa that experience consistent longitudinal shifts across multiple geographic regions and growing seasons [17]. Previously, we showed that drought-stressed rice root communities are developmentally delayed compared to well-watered communities [17]. Similarly, a study in sorghum reported a nearly complete halt in microbiota turnover during pre-flowering drought stress [18]. Assessing the impact of this developmental delay in the recovery period can reveal the extent to which drought disrupts the temporally coordinated interplay between host and root microorganisms.

As drought episodes become longer and more frequent [1, 21], it is necessary to determine the impact of an increasingly changing environment on plant-associated microbiomes. In particular, evaluating the resilience of root communities (*i.e*., their rate of recovery after a disturbance) can help us determine the permanence of drought-mediated alterations through the life cycle of their host. As highlighted in a recent review, there is a “need for improved mechanistic understanding of the complex feedbacks between plants and microbes during, and particularly after, drought” [8]. Here, we present a detailed temporal profiling of the rhizosphere and endosphere communities of rice plants grown under a range of drought stress durations. We find that extended drought produces lasting changes to root microbiota composition, with persistent phyla-dependent patterns of enrichment and depletion, and involving putative beneficial microbes. The findings have implications relevant to strategies to harness microbial communities for drought-tolerance in field crops.

## Results

### Experimental design and sequencing stats

To characterize the effect of drought on the temporal progressions of root-associated communities, we exposed rice plants (*Oryza sativa* ssp. *japonica* variety M206), grown in agricultural soil under controlled greenhouse conditions, to one of three increasingly longer drought periods: DS1 (11 days), DS2 (21 days), and DS3 (33 days). Given that microbiome succession is highly dynamic during the vegetative growth phase of rice [17], all drought treatments were initiated at 41 days after transplantation, before plants transitioned to the reproductive stage and microbiome composition stabilized. As a control treatment (WC), we kept an additional set of rice plants under well-watered conditions throughout the whole experiment. For each of the four watering regimes (WC, DS1, DS2, and DS3), plants were consistently sampled every ~10 days for a total of 13 collection time points spanning 136 days. This collection scheme covered the complete life cycle of rice and allowed us to track microbiome succession before, during, and after drought (**Figure 1A**). For each plant sampled, we profiled the bacterial and archaeal diversity associated with the rhizospheric and endospheric communities via high-throughput amplicon sequencing of the V4 region of the 16S rRNA gene. After filtering organellar sequences and removing non-persistent OTUs (defined as OTUs not present in at least 5% of all samples), we identified 4,135 OTUs (mean sequencing depth = 20,740 reads).

**Figure 1.**
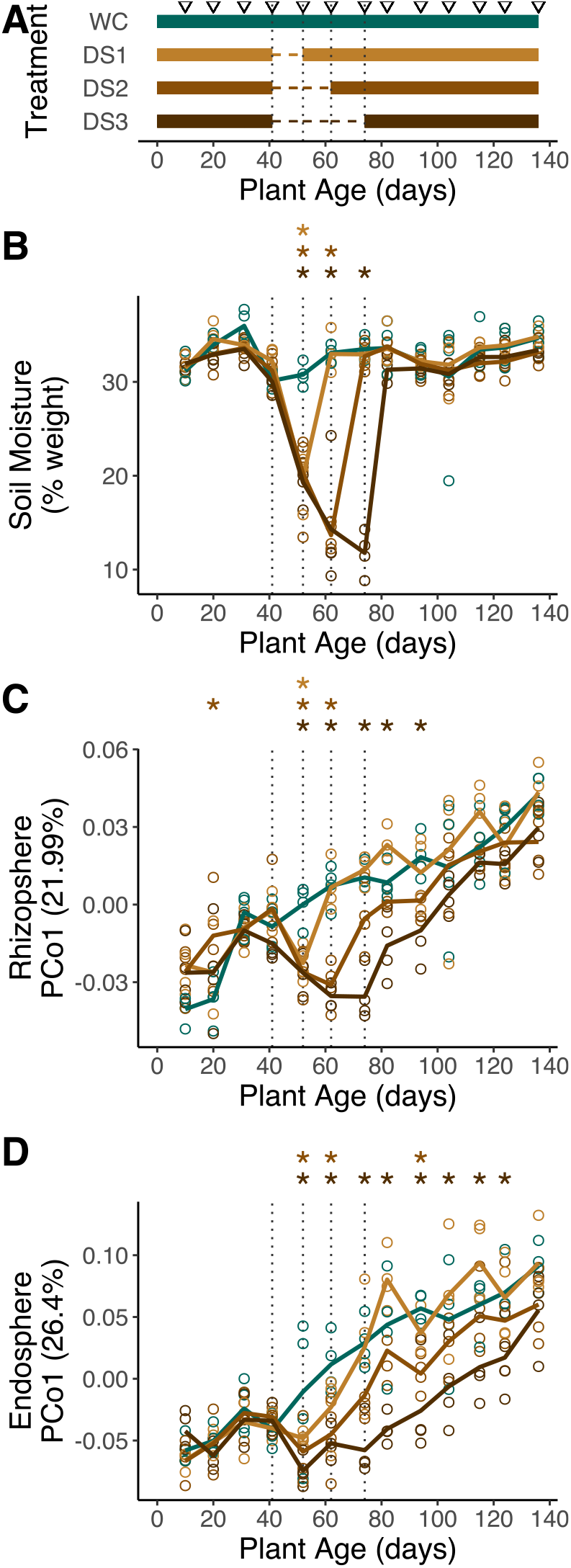
Compositional dynamics of rhizosphere and endosphere communities before, during, and after drought. **(A)** Timeline of the watering regimes followed by control (WC) and drought-treated (DS1, DS2, and DS3) plants. Horizontal lines represent the watering status during the experiment: solid segments indicate periods of constant irrigation while dotted segments indicate periods of suspended irrigation. Upside down triangles mark each of 13 collection time points spanning the complete life cycle of rice plants. **(B)** Soil percent moisture as measured by gravimetric water content **(C**, **D)** Beta-diversity patterns in the rhizosphere **(C)** and endosphere **(D)** communities. In both cases, the y-axis displays the position of each sample across the first principal coordinate (PCo) from a weighted UniFrac PCo analysis and the x-axis displays the age of the plant at the moment of sample collection. The trend lines in panels **B**, **C**, and **D** represent the mean values for each treatment throughout the experiment; asterisks on top indicate a significant difference (ANOVA, adjusted P < 0.05) between the control and each of the drought treatments at a specific time point.

### Beta-diversity patterns

Root compartment was the major driver of microbiome composition as evidenced by a clear separation between rhizosphere and endsophere communities across the first axis of an unconstrained principal coordinates analysis (PCoA) performed on weighted UniFrac distances (**SFigure 1A**). Moreover, a permutational multivariate analysis of variance (PerMANOVA) indicated that root compartment explained more than 62.8% of the variation in the whole dataset (P < 0.001). Therefore, to better explore the impact of drought treatment, collection time, and their interaction on each compartment, we ran a PerMANOVA on rhizosphere and endosphere samples independently. In both cases, all main and interaction effects were significant (**Table 1**).

**Table 1.** Influence of experimental factors and their interaction on the beta-diversities of rhizosphere and endosphere communities.

|  | Rhizosphere |  | Endosphere |  |
| --- | --- | --- | --- | --- |
| | $R^2$ | $P$ | $R^2$ | $P$ |
| Time | 0.1491 | 0.001 | 0.1710 | 0.001 |
| Treatment | 0.0459 | 0.001 | 0.0592 | 0.001 |
| Time x Treatment | 0.0171 | 0.047 | 0.0240 | 0.004 |
| Residuals | 0.7879 |  | 0.7457 |  |

Permutational multivariate analyses of variance were performed on weighted UniFrac distances.

To further examine the effect of drought duration on microbiome dynamics, we explored the longitudinal trends of beta-diversity captured by the first axis of independent PCoAs performed on each compartment (**Figure 1C-D, SFigure 1C-F**). In both rhizosphere and endosphere communities, PCo1 tracked the compositional development that root communities undergo during the lifecycle of rice plants as evidenced by the progressive transition of early to late time points along the axis. Additionally, PCo1 displayed drought-mediated shifts in community composition throughout time: while all watering regimes followed similar trajectories before drought onset (41-day-old mark), drought-treated plants started diverging from well-watered communities as soon as irrigation was suspended. The separation between control and stressed communities increased for as long as drought conditions were kept, with 31-day-stressed communities (DS3) showing the largest deviation from well-watered samples. Finally, drought treatments presented differential recovery dynamics upon rewatering: while both DS1 and DS2 samples recovered relatively quickly, DS3 communities remained significantly altered after drought stress was ceased (adjusted P < 0.05, asterisks in **Figure 1C-D**). This significant deviation from controlled communities was sustained for 50 days in the endosphere whereas it only lasted for 20 days in the rhizosphere, suggesting potential differences in community resilience across compartments. Such pattern contrasts with the temporal trends observed in soil water content as soil percent moisture was significantly reduced for all drought treatments during the stress period but immediately returned to control levels after irrigation was resumed (**Figure 1B**). Thus, despite soil water content being fully restored, prolonged drought hinders the ability of root communities to quickly recover.

### Drought-responsive taxa follow distinct longitudinal trends within and between compartments

To identify taxa affected by watering regime throughout time, we fitted negative binomial models to the relative abundances of individual OTUs and ran pairwise Wald tests contrasting well-watered controls (WC) against each drought treatment (DS1, DS2, and DS3) in each compartment at each collection time point. We found a total of 428 rhizospheric OTUs and 284 endospheric OTUs affected by treatment in at least one comparison (**Figure 2A, STable 1**, adjusted P < 0.05). The temporal distribution of significant effects among these differentially abundant OTUs followed distinct patterns in each compartment: in the rhizosphere, significance was mostly observed during the drought period; in the endosphere, it widely extended to the recovery phase of the experiment, especially for treatment DS3.

**Figure 2.**
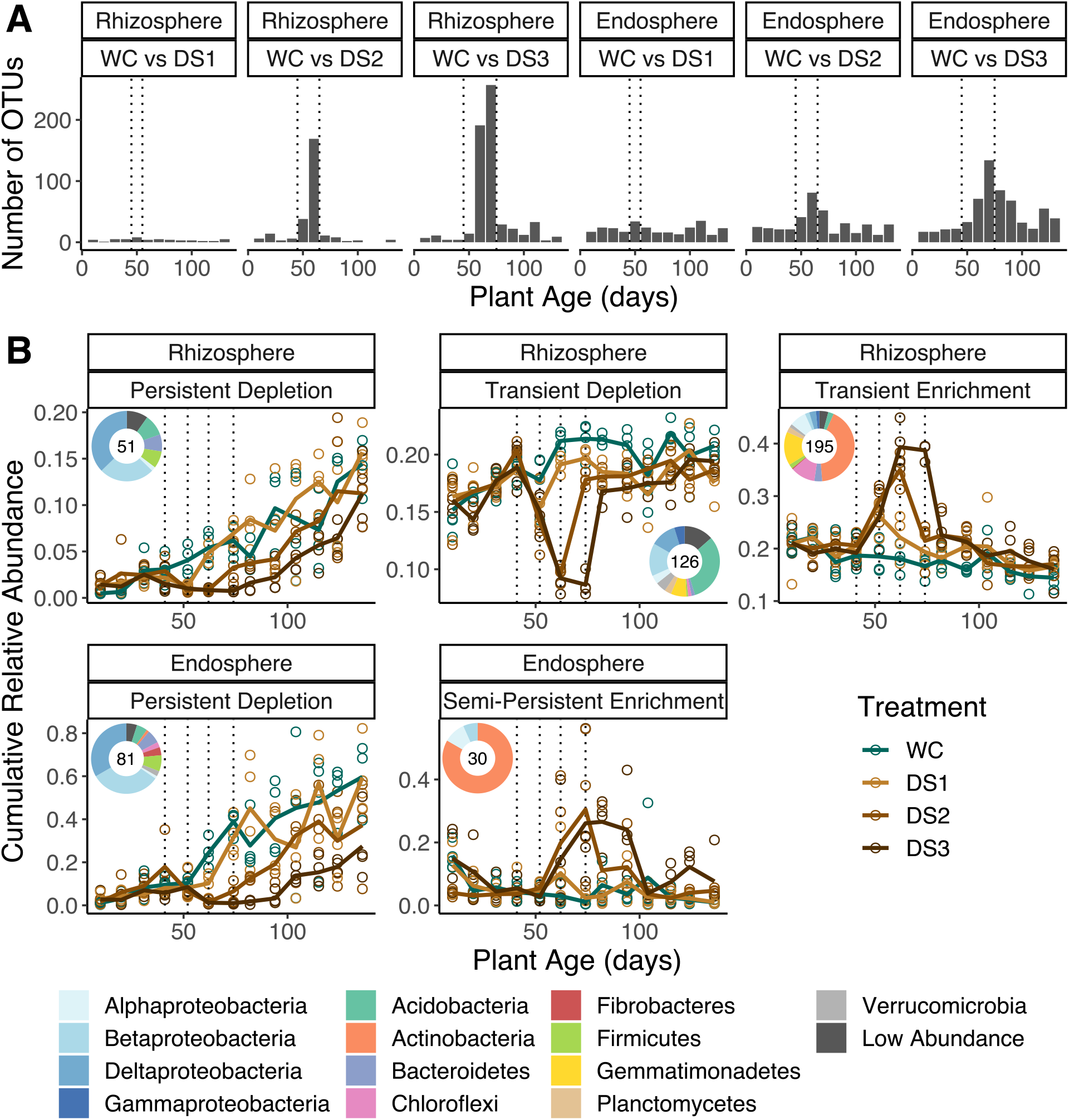
Drought-responsive OTUs show distinct longitudinal trends within and between compartments. **(A)** Number of differentially abundant OTUs (Wald test, adjusted P < 0.05) detected between well-watered controls and each of the drought treatments at each timepoint. **(B)** Longitudinal shifts in the cumulative relative abundances of rhizosphere and endosphere drought-responsive modules detected through hierarchical clustering. The complete set of detected clusters is shown in Supplementary Figures 2-3. Trend lines represent the mean values for each treatment throughout the experiment and inset donut plots display the size (number of OTUs) and taxonomic composition of each module. In all panels, the vertical dotted lines delimit the periods of suspended irrigation for each of the drought treatments.

While this approach detected clear ecological signals driven by drought stress (*e.g*., the number of differentially abundant OTUs was proportional to duration of stress), it also identified OTUs affected by other, potentially stochastic, processes. For example, multiple OTUs were found to be significantly affected by watering treatment in the collection time points preceding drought onset, when conditions were identical across treatments (**Figure 2A**). This effect was more pronounced in the endosphere communities, which exhibited greater within-group variation than rhizosphere communities (**SFigure 1B**). Thus, to identify coherent patterns of drought response in the set of differentially abundant OTUs, we performed hierarchical clustering on the log_2_ fold changes computed across all comparisons. (**STable 2, SFigures 2**). This method distinguished 3 rhizospheric and 2 endospheric modules displaying clear longitudinal trends across drought treatments. (**Figure 2B**).

One rhizospheric module consisted of 195 OTUs whose relative abundances increased under drought stress. Such enrichment was proportional to the duration of stress and was mostly constrained to the span of suspended irrigation in each treatment. The OTUs exhibiting this transient enrichment belonged mainly to the phyla Actinobacteria, Gemmatimonadetes, and Chloroflexi. In contrast, the other two rhizospheric modules showed clear signatures of abundance depletion under drought conditions, although each with unique recovery dynamics: while 126 OTUs were transiently depleted, *i.e*., their relative abundances were quickly restored after irrigation was resumed; 51 OTUs were persistently depleted, *i.e*., their relative abundances remained decreased weeks after stress was ceased. This latter pattern was particularly conspicuous in rhizospheres of plants that underwent 31 days of drought (DS3). While both depletion modules were enriched in OTUs classified as Acidobacteria, Betaproteobacteria, and Deltaproteobacteria, each one featured unique patterns at a lower taxonomic resolution (**SFigure 3**). On one hand, the majority of transiently depleted Betaproteobacteria belonged to order MND1 whereas almost all persistently depleted were Rhodocyclales. On the other hand, Deltaproteobacteria classified as Myxococcales and Desulfuromonadales were prominent in the transient and persistent modules, respectively.

Out of the 2 endospheric modules, one encompassed 30 OTUs enriched under drought while the other one contained 81 OTUs depleted under drought. In both cases, these shifts in abundances persisted after drought stress was suspended, albeit to different extents for each module. For OTUs positively impacted by drought, the increase in relative abundances lingered, depending on the specific treatment, up to ~10-20 days after irrigation was resumed. Interestingly, more than 80% of OTUs in this semi-persistently enriched module belonged to the phylum Actinobacteria. In contrast, for OTUs negatively impacted by drought, depletion relative to well-watered controls was observed throughout the whole recovery phase. Moreover, similar to the results observed in the rhizosphere, several persistently depleted OTUs were classified as Myxococcales and Rhodocyclales (**SFigure 3**). Together, these results indicate that phylogenetically distinct groupings of bacterial taxa follow diverse trajectories throughout drought stress and recovery in root-associated compartments.

### A highly occurring *Streptomyces* becomes the most abundant taxa in endosphere communities during and immediately after drought

Given the strong taxonomic signature displayed by the set of semi-persistently drought-enriched OTUs (**Figure 2B**), we further explored the compositional trends of each individual actinobacteria within this module. In particular, we calculated the abundance-occupancy curves of rhizosphere and endosphere communities and located each OTU along these spectra (**Figure 3A**). Overall, semi-persistently enriched actinobacteria were among the most abundant and occurrent members of root-associated communities, especially in the endosphere. One *Streptomyces* taxon, OTU 1037355, was notably predominant: not only was it detected in all collected samples, but its mean relative abundance was greater than that of 99% and 97% of all OTUs in the endosphere and rhizosphere communities, respectively. Furthermore, analyzing its temporal dynamics across treatments, we found that OTU 1037355 became the most abundant taxon in endosphere communities by the end of the DS2 and DS3 drought periods, reaching a mean relative abundance of 13.5% (**Figure 3B**). Additionally, OTU 1037355 remained the most abundant taxon in the endosphere during the early stages of recovery. In rhizosphere communities, the drought-mediated enrichment of OTU 1037355 was less prominent as it only reached a maximum relative abundance of 1.3% in drought-stressed samples. Moreover, even though the abundance of this OTU increased during the drought period, it immediately declined after irrigation was resumed. Thus, despite being significantly affected by drought in both communities, OTU 1037355 exhibited compartment-specific recovery trends.

**Figure 3.**
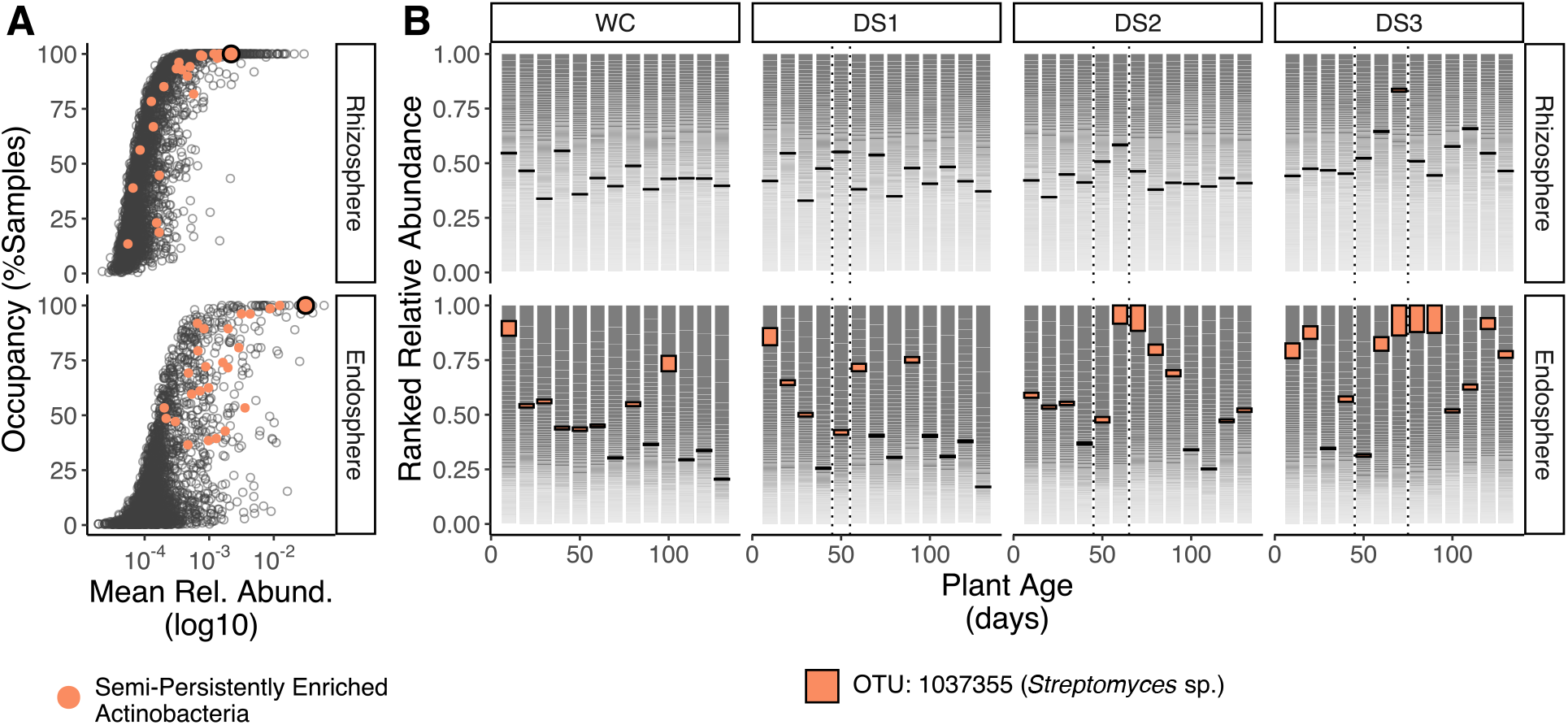
A drought-enriched OTU becomes the most abundant member of the endosphere communities. **(A)** Occupancyabundance curves for the rhizosphere and endosphere communities. The x-axis displays the log-transformed mean relative abundance of each OTU while the y-axis displays the percent of samples in which each OTU was detected. Actinobacteria OTUs detected as semi-persistently enriched in the endosphere (Fig 2B) are colored in orange and OTU 1037355 is further highlighted by a black outline. **(B)** Ranked relative abundances of individual community members throughout time. Each stacked bar plot displays all the OTUs detected in a particular time point: the height of individual bars represent the mean relative abundance of each OTU while the bar position across the y-axis indicates its rank within the community. The most abundant member of the semi-persistent enrichment module, *Streptomyces* sp. (OTU ID: 1037355), is highlighted. In all panels, the vertical dotted lines delimit the periods of suspended irrigation in each of the drought treatments.

We then assessed if the pattern of sustained enrichment displayed by OTU 1037355 was a reproducible feature of endosphere communities by analyzing its longitudinal dynamics in an independent drought experiment performed on the same rice cultivar grown in the same agricultural soil. Briefly, one-month old plants were drought-stressed for 21 days and allowed to recover for 7 days (see Methods). Samples were collected each week to track the drought-mediated temporal shifts and post-disturbance trends. The microbial profiles confirmed that this OTU was significantly enriched during and immediately after the imposition of drought conditions (**SFigure 4**). Moreover, this shift was even more conspicuous as the mean relative abundance of OTU 1037355 reached up to 24.0% of the total community in drought-stressed samples.

### A *Streptomyces* isolate classified as OTU 1037355 is a root growth promoting bacteria

To assess if this highly occurring *Streptomyces* taxon was part of the readily culturable fraction of the root microbiota, we screened a set of bacterial isolates previously collected from rice-associated rhizosphere and endosphere communities (see Methods) and found nine isolates classified as OTU 1037355. We then compared these isolates against the most prevalent sequence variant that mapped to OTU 1037355 in our longitudinal drought experiment; this sequence variant comprised 63.4% of all sequences mapping to that OTU (**SFigure 5A**). Five of the isolates differed by a single nucleotide, and one isolate, SLBN-177, was additionally derived from the same soil source as the longitudinal drought experiment. The full 16S rRNA gene of SLBN-177 was sequenced, and compared against the NCBI 16S rRNA gene database to further refine its taxonomic classification. We found that SLBN-177 shared 100% similarity with sequences from *Streptomyces pratensis, Streptomyces anulatus*, and *Streptomyces praecox (S. praecox* has been proposed as a synonym of *S. anulatus* [22]). Due to the sequence similarity and source of isolate, we further investigated the effects of SLBN-177 on rice phenotypes.

To evaluate the effect of *Streptomyces* SLBN-177 on rice growth phenotypes, seeds were inoculated with one of three microbial treatments: SLBN-177, SLBN-111, or a mock control. SLBN-111 is an Actinobacteria isolate from the genus *Microbacterium* and its associated OTU, 1108350, was found in low abundance in both the rhizosphere and endosphere communities (**SFigure 5B**). Unlike many other Actinobacteria taxa, OTU 1108350 was not significantly altered by drought in any compartment. Due to the weak association of OTU 1108350 with the plant and its stability under drought, SLBN-111 was selected to distinguish the effects of SLBN-177 on rice seedlings from a general response caused by the introduction of a high abundance of a foreign microbe. Inoculated seeds were grown for 10 days in an axenic closed system, followed by a 14 day period of non-sterile drought stress in an open system (with half the plants still fully watered), followed by 7 days of recovery. Plants were then harvested and root and leaf growth parameters were measured (**Figure 4A, SFigure 6**). A principal component analysis revealed that both watering and microbial treatments influenced the phenotypes of rice plants (**Figure 4B**). Watering treatment was the driving factor separating samples along the first axis while microbial treatment distinguished samples along the second axis. Interestingly, SLBN-177-inoculated plants clustered separately from mock- and SLBN-111-inoculated plants. Furthermore, root length was the main variable distinguishing microbial treatments (**Figure 4B**; **STable 3**). Notably, contrasts demonstrated that roots of SLBN-177-treated plants were significantly longer than mock- and SLBN111-treated plants in both well-watered and drought conditions (**Figure 4C**). Microbial treatments did not significantly affect any other measured trait; however, all phenotypic measurements were significantly reduced by drought (**SFigure 7A, STable 3**).

**Figure 4.**
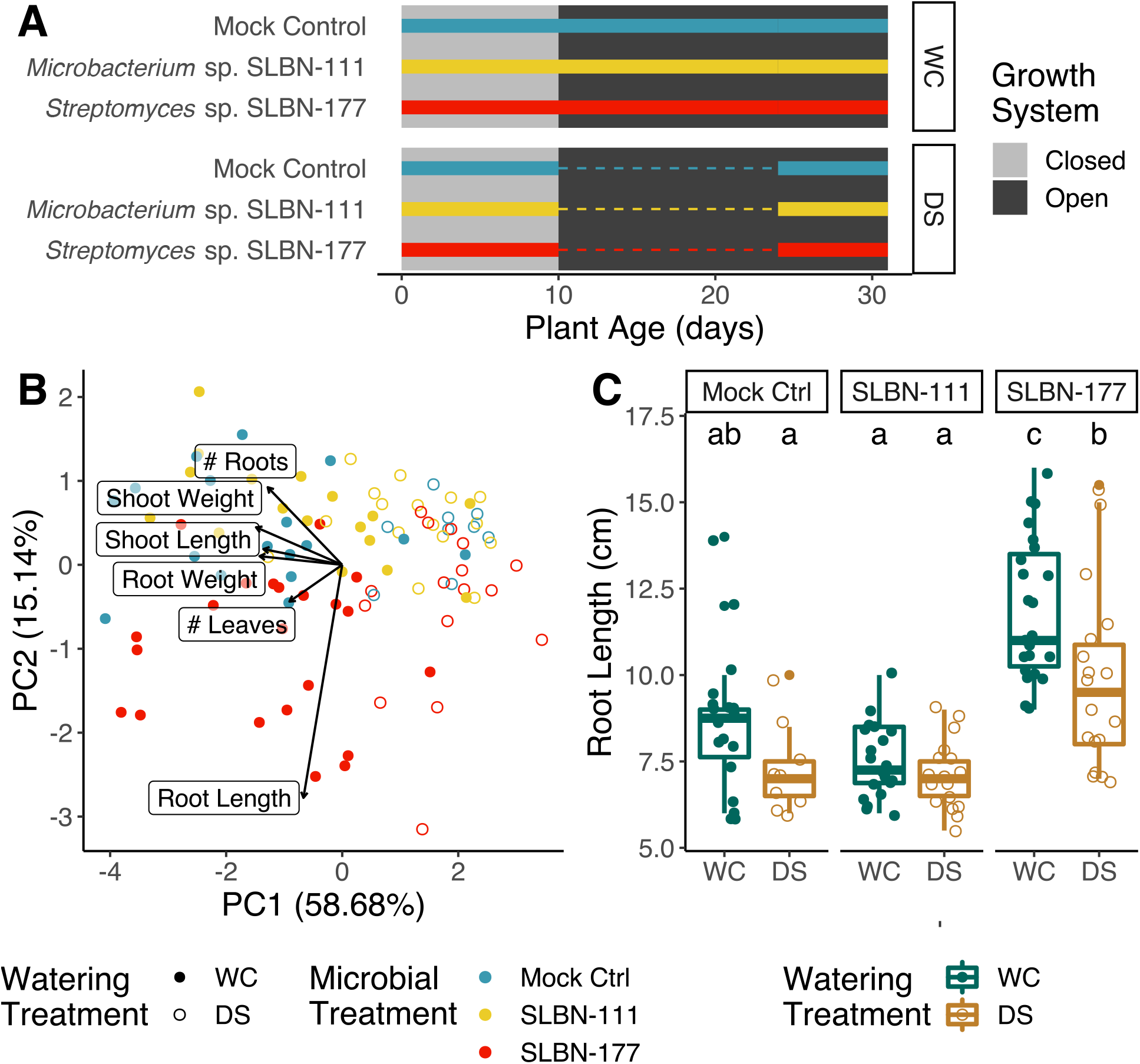
*Streptomyces* sp. SLBN-177 significantly increases root length under controlled conditions. **(A)** Timeline of the watering regimes followed by control (WC) and drought-treated (DS) plants. Horizontal lines represent the watering status during the experiment: solid segments indicate periods of constant irrigation while dotted segments indicate periods of suspended irrigation. Colors indicate the microbial treatment applied at the beginning of the experiment. The background colors indicate the periods during which the plant growth system was closed (*i.e*. axenic) or open. **(B)** Principal component analysis of plant phenotypes. Points represent individual plants harvested at the end of the experiment and vectors indicate the contribution of each of the measured variables. The microbial treatment and watering regime received by each plant are indicated by color and shape, respectively. **(C)** Distribution of root lengths across microbial treatments and watering regimes. Letters at the top indicate significantly different groupings (Tukey test, adjusted P < 0.05).

To explore potential mechanisms responsible for the root elongation, the genome of SLBN-177 was sequenced, assembled, and annotated. The assembly yielded 7.78 MB of sequence and 6,975 putative coding sequences. Mapping genes to KEGG pathways identified genes involved in the production of indole-3-acetic acid (IAA) through the indole-3-acetamide (IAM) pathway, including *iaaM* (a tryptophan 2-monooxygenase) and *amiA2* (a putative amidase), potentially involved in the first and second steps of this pathway, respectively. The *iaaM* gene shared an 88.5% amino acid similarity with homologs from *Streptomyces coelicolor* and 88.9% with homologs from *Streptomyces scabiei*, both of which have previously been implicated in IAA biosynthesis [23, 24]. Additionally, we identified gene clusters associated with siderophore and antimicrobial biosynthesis (**STable 4)**.

To confirm that SLBN-177 colonized the roots of rice plants, we performed 16S rRNA gene profiling on the endospheres of a subset of samples and compared the relative abundance of microbial reads to organellar reads. The mean relative abundance of OTU 1037335 on SLBN-177-treated plants reached 5.7% and 30.7% in well-watered and drought-recovered samples, respectively, suggesting the enrichment of SLBN-177 persists in the recovery phase, as observed in OTU 1037355 in the previously described experiments (**SFigure 7B**). In contrast, OTU 1108350 was barely detected in SLBN-111 treated samples, reaching a maximum relative abundance of 0.001%. Two drought-recovered control plants also had notable relative abundances of OTU 1037355, which could be a consequence of the open system portion of the experiment. However, the relative abundances of OTU 1037355 in these plants were much lower than drought-recovered plants inoculated with SLBN-177 (**SFigure 7B**). Collectively, these results indicate that OTU 1037355 is a plant-growth promoting *Streptomyces* that is a key contributor to the compositional dynamics of endosphere communities during drought and recovery.

### Drought permanently delays rhizosphere and endosphere microbiome development

Relative abundances of root-associated taxa follow reproducible longitudinal trends that can be used to track root microbiome maturation throughout time by training random forests models [17]. Using this approach on field-grown samples, we have previously shown that drought-stressed plants host a developmentally immature microbiota [17]. Given this result, however, it is unknown whether microbiome immaturity persists upon rewatering. To explore this possibility, we used samples from well-watered plants to train seperate full random forest models for each compartment by regressing OTU relative abundances as a function of host chronological age. For each compartment, we ranked each OTU based on age-predicting importance and selected the top 65 (a threshold identified through cross-validation - **SFigure 8A**) to generate sparse random forest models (**STable 4**). Similarly to the age-discriminant taxa detected in our previous field study [17], these top OTUs could be classified as early, late, or complex root colonizers based on their relative abundance patterns through time: early colonizers displayed initial high abundances that progressively declined, late colonizers exhibited initial low abundances that progressively increased, and complex colonizers comprised OTUs that didn’t fit any of these two trends (**SFigure 8C, STable 5**). Among the set of early endosphere colonizers, most were classified as Chloroflexi and Betaproteobacteria (mainly Burkholderiales), whereas the set of early rhizosphere colonizers were more phylogenetically diverse. In contrast, both compartments had a clear enrichment of Deltaproteobacteria (mainly Myxococcales) and Betaproteobacteria (mainly Rhodocyclales) in the set of late colonizers (**SFigure 9**).

The 65-taxon sparse models explained the 89.06% and 90.08% of variance related to plant age in the rhizosphere and endosphere communities, respectively. Furthermore, these models accurately predicted plant age on a validation set of well-watered samples, indicating that this approach was able to capture the consistent taxonomic shifts observed during normal root microbiome succession. We then applied the sparse random forest models to each of three drought regimes to assess the effect of drought on microbiome succession. We observed a clear deviation from the baseline development established by well-watered controls (**Figure 5A**). To further measure this divergence, we calculated the relative microbiome maturity of each sample as the difference between the predicted microbiome age and the baseline microbiome age of well-watered plants collected at the same chronological age (**Figure 5B**). The results showed that, before drought onset, all watering regimes tracked normal microbiome development. However, microbiome progression was interrupted during drought and relative microbiome maturity became increasingly delayed. Furthermore, the extent of this microbiome immaturity was proportional to the duration of stress, with DS3 communities showing the highest departure from baseline development. For DS2 and DS3 samples, this microbiome immaturity persisted throughout the rest of the life cycle, even after irrigation was resumed.

**Figure 5.**
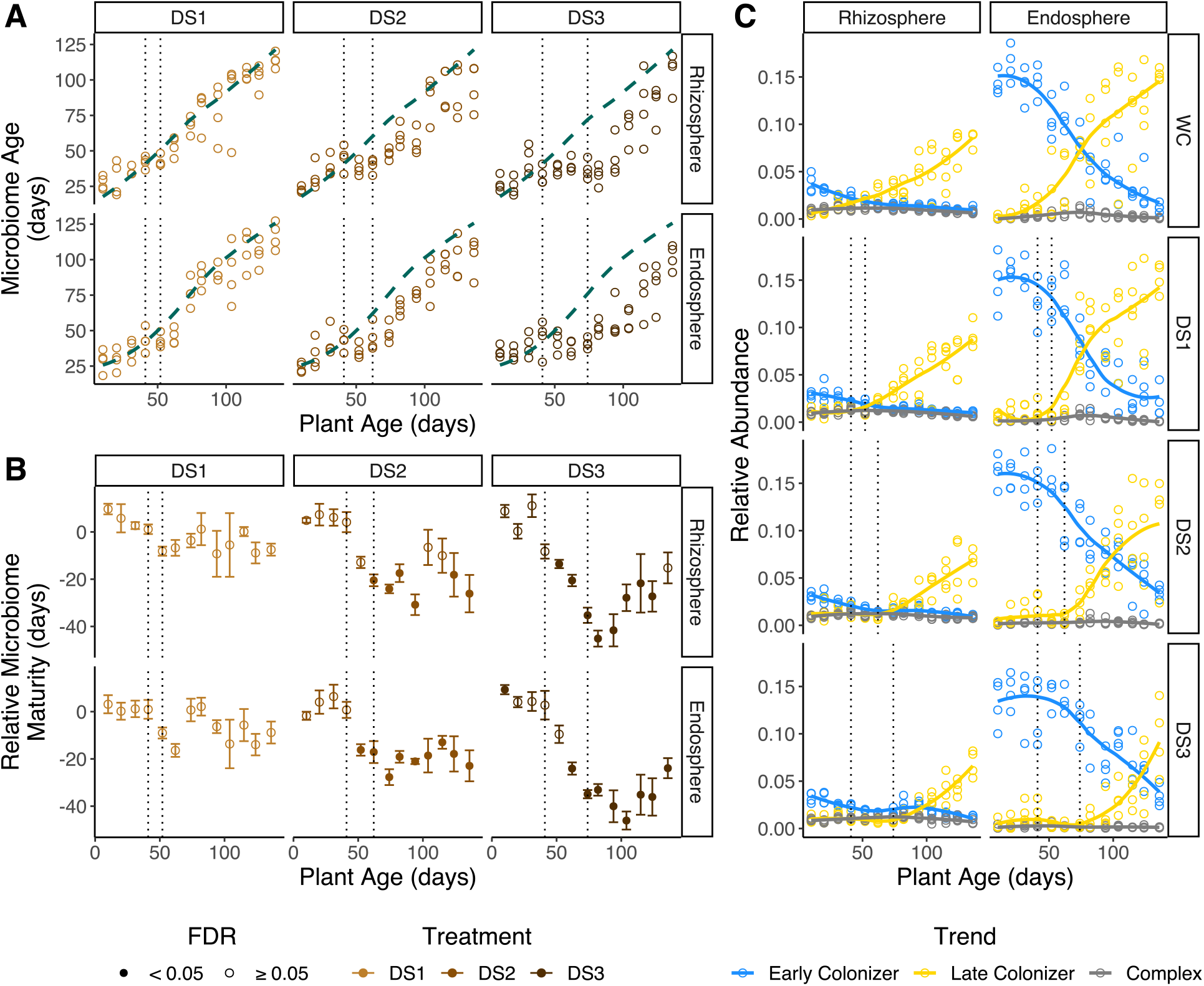
Persistent immaturity of root microbiomes in drought-stressed plants. **(A)** Microbiome age predictions of rhizosphere and endosphere communities across drought treatments (D1, D2, and D3). The dashed curve represents the baseline microbiome development under well-irrigated conditions and was calculated by fitting a smoothed spline between the predicted microbiome age and the chronological plant age in the control (WC) test set. **(B)** Relative microbiome maturity measured as the difference between the predicted microbiome age and the interpolated value of the smoothed spline at each sampling time point. Solid points indicate a significant difference (ANOVA, adjusted P < 0.05) between the control and each of the drought treatments. **(C)** Longitudinal shifts in the aggregated relative abundances of the early, late, and complex colonizers used in the random-forest models.

To understand the compositional changes driving the drought-mediated delay in root microbiome development, we analyzed the abundance patterns of age-discriminant taxa across watering treatments. In both compartments, we observed a clear shift in the transition of dominance between early and late colonizers (**Figure 5C**). In the rhizosphere, this transition was detected at the ~50 and ~90 day-old marks in WC and DS3 plants, respectively; in the endosphere, the transition was detected at the ~70 and ~120 day-old marks in WC and DS3 plants, respectively. This temporal shift in root microbiome assembly was mostly linked to a delay in the onset of late colonizers as evidenced by a persistent decrease in their relative abundances upon drought stress. Additionally, there was a considerable overlap between the set of late colonizers and the differentially abundant OTUs assigned to the persistently depleted modules detected in the rhizosphere and endosphere communities (**SFigure 8B**). Overall, these results indicate that drought stress permanently delayed microbiome development by affecting the recruitment of late colonizers.

## Discussion

Here, we provide a detailed characterization of the drought-mediated changes and post-disturbance dynamics of root microbiomes through the life cycle of rice plants. We found that both the magnitude of compositional changes undergone during drought and the capacity to fully recover upon rewatering were significantly affected by the duration of drought stress experienced by the host and its associated bacterial and archaeal communities. In particular, we show that prolonged drought led to a severe microbiome restructuring that persisted even after irrigation was reestablished. Moreover, endosphere communities remained altered longer than rhizosphere communities, suggesting that the compartment-specific responses to drought previously reported for rice [12] and other angiosperms [13, 14] extend to the recovery period. The observed permanence of drought-mediated changes contrasts with the rapid resilience recently reported for the root microbiomes of sorghum, in which the relative abundances of Actinobacteria progressively increased over the course of six weeks of drought but quickly returned to pre-drought levels within a week of rewatering [18]. These conflicting results could stem from differences in drought tolerance between the two crops since sorghum, a naturally drought tolerant C4 plant with a deep root system, is better adapted to arid conditions than rice, a C3 plant with shallow roots [25–27]. Expanding the characterization of recovery dynamics to a broader range of hosts is necessary to advance our understanding of root microbiome resilience.

Hierarchical clustering of differentially abundant OTUs further revealed modules of drought-responsive taxa with distinct recovery dynamics: while some OTUs were transiently impacted by drought (*i.e*., changes in their relative abundances were mainly constrained to the period of stress), others were persistently affected throughout recovery. These diverse recovery patterns could stem from differences in life strategies and metabolic capabilities among root-associated microorganisms. For instance, copiotrophs could recover more quickly than oligotrophs as they exhibit higher growth rates and lower resource use efficiency [28]. Given that rewetting of dry soils releases specific forms of carbon and nitrogen to the environment [29], the ability to metabolize these liberated resources could also facilitate a quick recovery. For example, soil bacteria communities recovering from drought have been associated with CO2 and N2O fluxes [30]. In contrast, the re-oxygenation of soils during drought could impact anaerobic microorganisms. Under flooded conditions, rice paddies become more reduced over time, allowing a succession of taxa that respire increasingly energetically unfavorable compounds [31]. Events that re-oxygenate the soil inhibit growth and activity of anaerobic bacteria that require a reduced state for more energetically unfavorable forms of metabolism [32, 33]. Notably, several persistently depleted taxa are known anaerobes that could be affected negatively by re-oxygenation, including genera Desulfovibrio (reduces sulfate), Geobacter (reduces iron and other metals), and Anaeromyxobacter (reduces various metals)[34–37]. Future metagenomic surveys of the functions enriched across different recovery strategies could help us better understand the processes driving this temporal differentiation.

Additionally, plant-microbe interactions could impact the responses of rhizosphere and endosphere microorganisms during and after drought. Root exudation is a temporally dynamic process that can promote or inhibit the growth of particular microbial taxa [38]. It has been shown that drought and rewetting can modify exudate composition [39–41], which in turn could alter the activity of microorganisms at the root-soil interface [42]. Drought can also affect the growth and architectural properties of roots [43], potentially reshaping microbiome composition [44, 45]. Finally, rice plants can respond to drought stress by delaying flowering anywhere from 4.5 days to 22 days [46–49]. This developmental arrest could impact root microbiome assembly processes that rely on temporally-staged host-mediated signaling. Interestingly, using a random forest approach, we found that the temporal progressions of rhizosphere and endosphere communities were also interrupted during prolonged drought stress (DS2 and DS3). While similar delays in microbiome development have been recently reported [17, 18], our post-disturbance sampling scheme allowed us to further evaluate if this drought-associated immaturity persisted after irrigation was resumed. We found that root communities remained underdeveloped throughout the whole recovery period due to a delay in the arrival of late root colonizers, many of which were part of the OTUs identified as persistently depleted in our differential abundance analysis. Late colonizing taxa have been recently shown to follow reproducible temporal abundance patterns in the rhizosphere and endosphere communities of different rice genotypes grown across geographically distant areas and over multiple growing seasons [17]. This high degree of conservation suggests that plant selectivity might play a key role in the late-stage assembly dynamics of root microbiomes, further hinting that the drought-induced delay of late colonizers observed in our study might be linked to host-mediated processes.

A characteristic response of host-associated drought-stressed microbiomes is the relative enrichment of Actinobacteria, which is broadly shared across a wide diversity of plants [12–14] and has been shown to correspond to an increase in the absolute abundance of members of this phylum [18, 50]. While the implications of Actinobacteria enrichment are not fully understood, there is evidence that it is at least partially mediated by the host [18]. Furthermore, in the context of drought stress, a recent study found a positive correlation between drought tolerance in angiosperms and the relative abundance of an endospheric *Streptomyces*, suggesting a potential beneficial role under stress [14]. Here, we have found that, in addition to a drought-mediated increase in their relative abundances, several Actinobacteria displayed a unique recovery trend that is both compartment-specific and dependent on the degree of drought stress. The most prominent member of these taxa, OTU 1037355, became the most abundant member of the endosphere community during and immediately after the stress period. This pattern of sustained enrichment and prevalence was further confirmed in an independent drought experiment, indicating that this is a reproducible temporal trend of the root-associated communities of rice. Inoculation of rice seedlings with a corresponding isolate, SLBN-177, resulted in increased root length under both drought and well watered conditions. Genome sequencing of SLBN-177 identified the iaaM gene, encoding the enzyme Trp-2-monooxygenase involved in the first step of the auxin analog indole-3-acetic acid (IAA) through the indole-3-acetamide pathway [51]. This gene has previously been identified in plant-growth promoting Streptomyces [23]. Also identified in the genome was the putative amidase amiA2, which could catalyze the second reaction of indole-3-acetamide to IAA. While the increased root growth in our experiment suggests an Actinobacteria-mediated increase in phytohormone biosynthesis, our system was not designed to test other common methods of plant growth promotion employed by Streptomyces, specifically by protecting the host through the inhibition of opportunistic pathogens. The genome of SLBN-177 contains gene clusters involved in the synthesis of antibiotics, which could inhibit the activity of these pathogens, and siderophores, which can provide iron to the host as well as trigger induced systemic resistance [52–54]. The presence of these gene clusters suggests that a further investigation of SLBN-177 is needed to fully understand its interaction with the host plant.

The prominence of SLBN-177 in the microbial community during and after drought, paired with its plant growth promoting mechanism, suggests functional implications for the microbiome restructuring and persistence. Given that highly abundant community members are likely to play key roles in the network of microbe-microbe and host-microbe interactions, it is possible that the enrichment of this OTU could have a major impact on rice plants during critical periods of environmental fluctuations. As extreme climate events become more prevalent, crops will likely experience multiple periods of intermittent drought within a growing season [55], and the ability to quickly recover and prepare for future drought events could be vital for survival. Plants are able to prepare for these future drought events through the development of a stress memory, a series of morphological, molecular, and physiological modifications that plants undergo during an initial drought episode that primes a more robust response to subsequent drought events [56, 57]. For example, some rice cultivars increase their root plasticity in the recovery period after an initial drought event, which allows the roots to penetrate hardened soil [57]. After a repeat drought event, cultivars with increased root growth responses to the initial drought event had an increase in root water uptake, stomatal conductance and shoot growth compared to those that did not [58]. As an extended root phenotype [59], rhizosphere and endosphere communities might also contribute to this stress memory. In particular, drought-mediated compositional changes that persist during the recovery period could amplify the response of plants to future drought events.

## Methods

### Experimental design

All data presented in this study were gathered from two controlled greenhouse experiments and one controlled growth chamber experiment performed at the University of California-Davis. The main study was carried out in the winter/spring of 2018, while the complementary study was a small pilot experiment carried out in the summer of 2017, and the growth chamber experiment was carried out in the winter of 2019.

#### Main Experiment

Twenty plastic containers holding 16 individual pots were arranged in a 5-by-4 configuration. Single seedlings were transplanted to each pot, and watering regimes were assigned to plastic containers in a randomized complete block design: each drought treatment (D1, D2, D3) was assigned to 4 blocks, while the well-watered treatment (WC) was assigned to 8 blocks (**SFigure 10**). The additional WC replicates were exclusively used to train the random forests models. Ten days after seedling transplantation, samples were collected every ~10 days (**Figure 1A**) for a total of 13 collection time points spanning 136 days. This design resulted in 4 biological replicates per treatment and collection time point combination.

#### Complementary experiment

Fifty potted plants were randomly assigned to one of two watering regimes: drought-stress treatment (DS) or well-watered controls (WC). Samples were collected at the 28, 35, 42, 49, and 56-day marks, encompassing 1 pre-drought, 3 drought, and 1 post-drought time points. This design resulted in 5 biological replicates per treatment and collection time point combination.

#### Semi-sterile phenotyping experiment

One hundred and fifty-six seeds were inoculated with either SLBN-177, SLBN-111, or a mock treatment in closed, sterile 75-ml culture tubes. After 10 days, half of the seedlings inoculated with each treatment began a two week period of drought stress followed by a week of recovery, at which point the plants were harvested.

### Plant growth

The rice variety used in this study was cultivar M206, an *Oryza sativa* subsp. *japonica* accession grown in California. Dehulled seeds were treated with a 50% bleach solution for 5 minutes followed by 5 washes with sterile water. Surface sterilized seeds were plated on Murashige and Skoog (MS) agar, and germinated in a growth chamber for 7 days. Seedlings were then transplanted to pots holding agricultural soil collected from a rice field in Arbuckle, California (39°0’42.235”N, 121 °55’19.632”W).

### Watering regimes

During non-drought periods, plants were irrigated *ad libitum* to keep the soil under submergence. Drought was imposed by draining all water from the plastic containers and allowing soils to dry. In the main experiment, DS1, DS2, and DS3 drought treatments started 41 days after transplantation and lasted for 11, 21, and 33 days, respectively (**SFigure 1B**). In the complementary study, drought started 28 days after transplantation and lasted for 21 days. At the end of the drought period, water was added to the plastic containers to recover the plants.

### Gravimetric water content measurements

For each pot collected, soil samples were harvested and placed in 15-ml Falcon tubes. After recording the initial weight, samples were allowed to dry inside a 42°C oven for 4 months. The dry weight of the samples was recorded and the percentage of moisture was calculated.

### Isolation of microbes

Bacterial colonies were isolated from rhizosphere and endosphere communities of rice plants derived from a previous study [12]. Briefly, rice plants were grown in three different agricultural soils (including the one used in this experiment) under controlled greenhouse conditions. One month-old plants were drought-stressed for three weeks and root systems were harvested. Isolates were then collected by plating both rhizosphere soil and ground root tissue resuspended in sterile phosphate-buffered solution on Actinomycete Isolation Agar (Himedia).

### Semi-sterile phenotyping experiment

Glass culture tubes (75 mL) were filled with 15 g wetted calcined clay. This setup was autoclaved twice for one hour with 24 hours between autoclave cycles. Rice seeds were sterilized by submerging seeds in 50% bleach for 15 minutes followed by 5 minutes of 70% EtOH, followed by five washes with sterilized H_2_O. Sterile seeds were placed in each tube. Isolate SLBN177 and SLBN111 were grown in LB liquid media and diluted in half-strength Murashige-Skoog media with no added sugar to an OD of 0.01. Ten mL of sterile MS or MS with one of the isolates were added to each seeded tube. Plants were grown in sterile conditions for 10 days. After this period, the culture tube lids were removed, and half the tubes were allowed to dry out for 14 days. The other half were watered periodically (2-3 days as needed) with sterilized H_2_O. After the drought period, all plants were well watered for a 7 day recovery period. Thereafter, plants were harvested, and shoot and root length and fresh weight were measured, as well as the number of leaves and roots. Sections of the roots were washed and flash frozen for 16S amplicon sequencing.

### Microbiome sample collection, processing, and DNA extraction

Root sample collection, compartment processing, and DNA extraction were performed as previously described [60]. Briefly, we scooped whole plants outside the pots and shook vigorously to remove all the soil not firmly attached to the roots. We then collected the 5 cm of root tissue immediately below the shoot-root junction in a 50-mL Falcon tube filled with 15 mL of sterile phosphate-buffered solution. Rhizosphere samples were collected by vortexing the roots and collecting 500 μl of the resulting soil suspension in PowerBead tubes (Mo Bio Laboratories). Endosphere samples were collected by washing the roots in fresh PBS to further discard any remaining soil and sonicating them three times (50 to 60 Hz for 30 s). Sonicated roots were placed in PowerBead tubes and homogenized by intense agitation for 1 min (Mini Beadbeater; BioSpec Products). DNA extractions were performed immediately after compartment separation, following the PowerSoil DNA isolation kit (Mo Bio Laboratories) protocol.

### 16S amplicon library preparation

Library construction followed a previously described dual-indexing strategy [60, 61]. For 16S rRNA gene libraries, the V4 region was amplified using the universal primers 515F and 806R. Amplification was carried out with the following touchdown PCR program: a first phase consisting of 95°C for 5 min, followed by 7 cycles of 95°C for 45 s, 65°C for 1 min (decreasing at 2°C/cycle), and 72°C for 90 s, with a second phase consisting of 30 cycles of 95°C for 45 s, 50°C for 30 s, and 72°C for 90 s, followed by a final extension at 72°C for 10 min. All PCR amplifications were performed using the HotStar HiFidelity polymerase kit (Qiagen). After running a 1% agarose gel to verify proper amplification, libraries were cleaned with AmPure XP magnetic beads (Beckman Coulter, Inc.), quantified (Qubit dsDNA HS assay kit; Thermo Fisher Scientific), and pooled in equimolar concentrations. Pooled libraries were then concentrated, gel purified (Nucleoscopic gel and PCR cleanup kit; Macherey-Nagel), quality checked (BioAnalyzer HS DNA kit; Agilent Technologies), and submitted for 2-by 250-bp Miseq sequencing (Illumina) to the DNA Technologies and Expression Analysis Cores at the UC Davis Genome Center (supported by NIH Shared Instrumentation Grant 1S10OD010786-01).

### 16S amplicon sequence processing

The paired-end reads were demultiplexed with custom scripts (https://github.com/bulksoil/BananaStand) and assembled into single sequences with PANDAseq [62]. Chimeric sequences were detected and discarded with usearch61[63]. OTU clustering at 97% identity was performed with the QIIME [64] implementation of UCLUST [63], using a close reference strategy against the 13_8 release of the Greengenes 16S sequence database [65]. OTUs classified as mitochondria and chloroplast were discarded from the OTU table (except in the semi-sterile phenotyping experiment), and non-prevalent OTUs (defined as OTUs not present in at least 5% of our samples) were filtered out.

### Genome sequencing

SLBN-177 was grown in liquid LB for 24 hours and DNA was extracted with Qiagen Blood and Tissue kit. DNA sequencing was done in the laboratory of Dr. Bart Weimer (UC Davis) as part of the 100K Pathogen Genome Project [66] as previously described [67–70]. Approximately 600 ng of purified gDNA was used to construct a sequencing library using KAPA HyperPlus library preparation kit (Roche Diagnostics). Final library QC for size distribution verification was done on Caliper Lab Chip ^GX^ (Perkin Elmer) and library quantification was done using KAPA Library Quantification Kit (Roche Diagnostics). Pooled libraries were sequenced on the Illumina HiSeq X Ten using a PE150 protocol. Reads were trimmed with Trimmomatic [71], assembled with SPAdes [72], and annotated with prokka [73], all with default settings. Contigs shorter than 1000 bp or with an average coverage less than 20X were excluded. KEGG Ontology terms were extracted from the prokka output using the script Prokka2KEGG (https://github.com/SilentGene/Bio-py/tree/master/prokka2kegg). Secondary metabolite biosynthesis gene clusters were identified using antiSMASH [74].

### Statistical analyses

All analyses were conducted in the R Environment version 3.5.1 [75]. For beta-diversity analyses, we used phyloseq [76] to calculate weighted UniFrac distances [77] on OTU counts normalized via variance-stabilizing transformation [78, 79]. Unconstrained principal-coordinate analysis was performed with the pcoa function from the ape package [80]. Permutational multivariate analyses of variance and canonical analyses of principal coordinates were performed with vegan [81]. Differential abundance analyses were performed with DESeq2 [78, 79]. Random forest modelling was performed using the randomForest package [82]. All plots were generated with ggplot2 [83].

## Data availability

Raw reads have been deposited in the SRA under Bioproject PRJNA551661.

## Code availability

All scripts and intermediate files have been deposited in GitHub (https://github.com/cmsantosm/RiceDroughtRecovery).

## Acknowledgments

This project was funded by the National Science Foundation (grant IOS 1444974) and the United States Department of Agriculture, Agricultural Experiment Station (project CA-D-XXX-6973-H). C.S.M. acknowledges support from the University of California Institute for Mexico (UCMEXUS), Consejo Nacional de Ciencia y Tecnología (CONACYT), and Secretaría de Educación Pública (México). C.S.M. and Z.L. also acknowledge partial support from the Elsie Taylor Stocking Memorial Research Fellowship and the Henry A. Jastro Graduate Research Award. J.E. is supported by USDA National Institute of Food and Agriculture Postdoctoral Fellowship [grant no. 2019-67012-2971/project accession no. 1019437]. We thank Dr. Ryan Melnyk for helpful advice regarding genome assembly, annotation, and interpretation.

## Author contributions

C.S.M., Z.L., and V.S. conceptualized the study; C.S.M, Z.L., J.E., and B.N. performed the experiments; B.H. and B.C.W. generated the SLBN177 genome sequence and reviewed manuscript; C.S.M and Z.L. analyzed the data; C.S.M., Z.L., J.E., and V.S. wrote the paper.

