## Supplementary Figures for "Drought duration determines the recovery dynamics of rice root microbiomes"

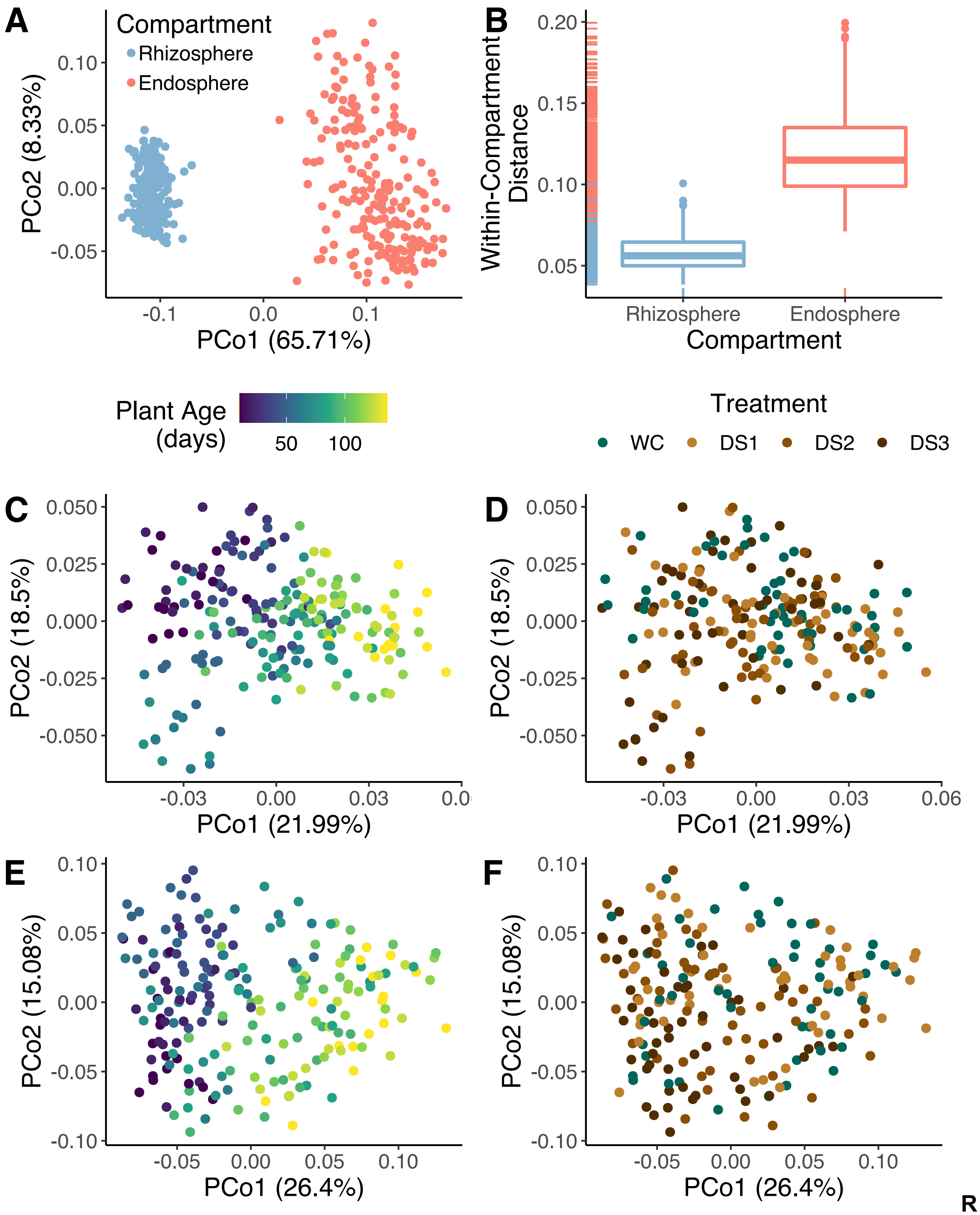


R

**Supplementary Figure 1.**

**Compartments harbor compositionally distinct microbial communities. (A)** Principal coordinate analysis (PCoA) performed on weighted UniFrac distances of the whole dataset. Colors indicate compartment. **(B)** Distribution of rhizosphere and endosphere within-group distances, *i.e.* distances between samples within each treatment and time point combination. **(C-F)** PCoA performed on the rhizosphere **(C-D)** and endosphere **(E-F)** subsets. Colors indicate plant age **(C & E)** and watering regime **(D & F)**

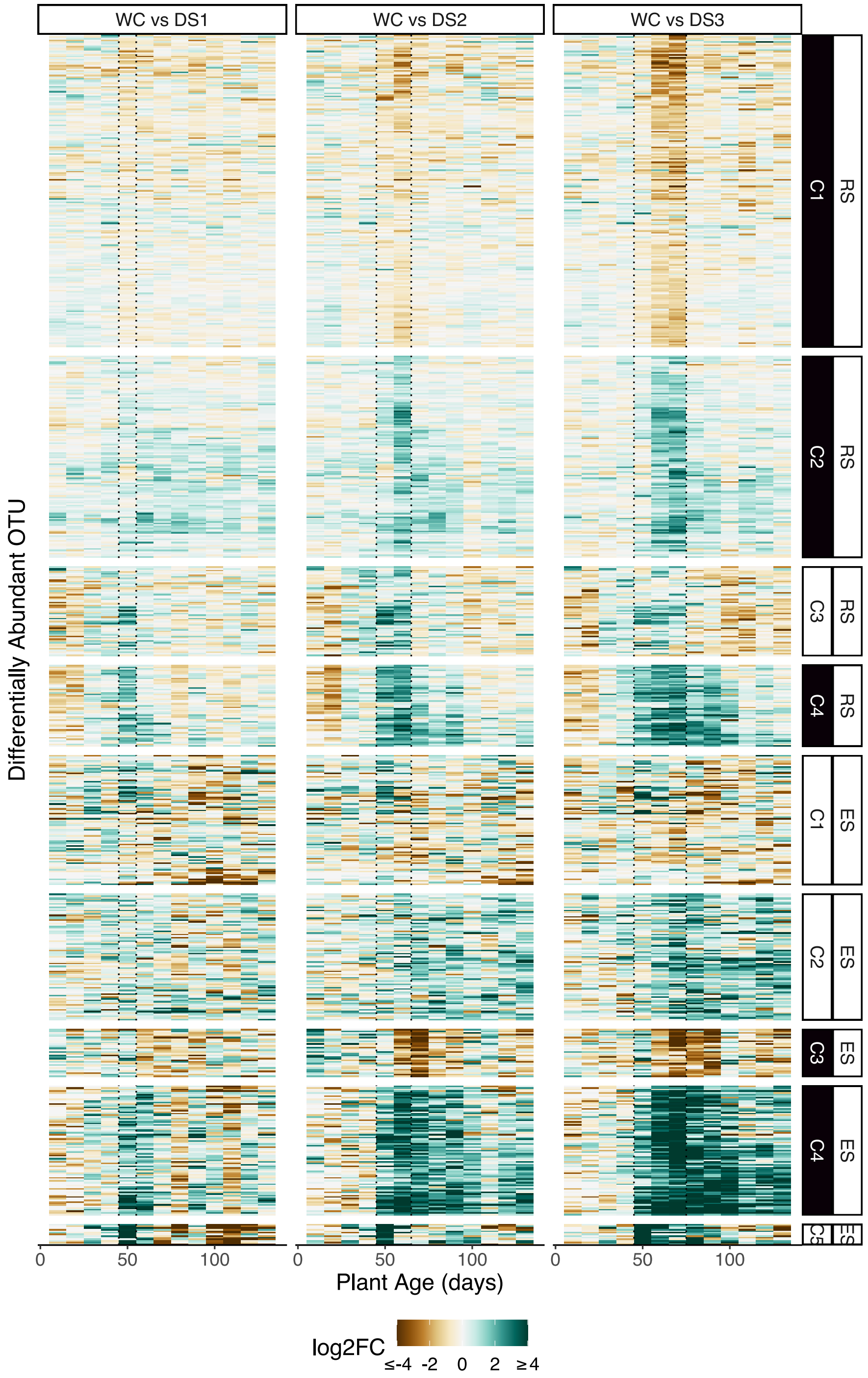


**Supplementary Figure 2.**

**Complete set of drought-responsive OTUs.** Heatmap displaying the log2 fold-changes between water controls and drought treatments: depletion under drought tends towards green while enrichment under drought tends toward brown**.** Each row represents a differentially abundant OTU detected as significant (Wald test, FDR < 0.05) in at least one pair-wise comparison. Horizontal facets indicate each of the modules detected through hierarchical clustering in the rhizosphere (RS) and endosphere (ES). Clusters shown in Figure 2B are highlighted in black. Vertical dotted lines delimit the periods of suspended irrigation for each of the drought treatments.

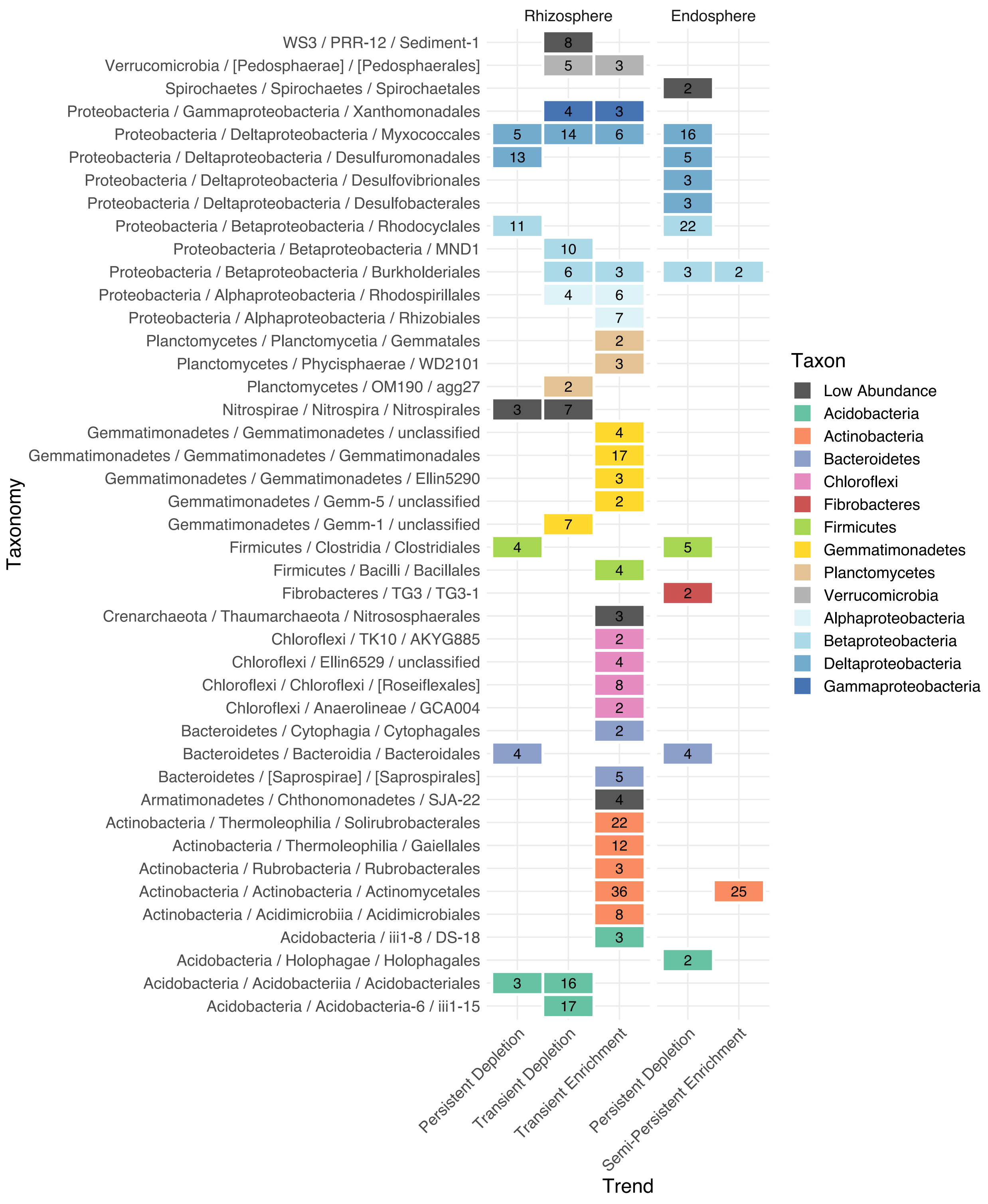


**Supplementary Figure 3.**
**Modules of drought responsive OTUs exhibit a strong taxonomic signature.** Classification of the differentially abundant OTUs in each of the distinct drought modules detected through hierarchical clustering. Each tile indicates the number of classified OTUs in a particular module, while the color represents membership to a specific Phylum / Proteobacteria class. Orders with only one representative have been excluded to ease visualization.

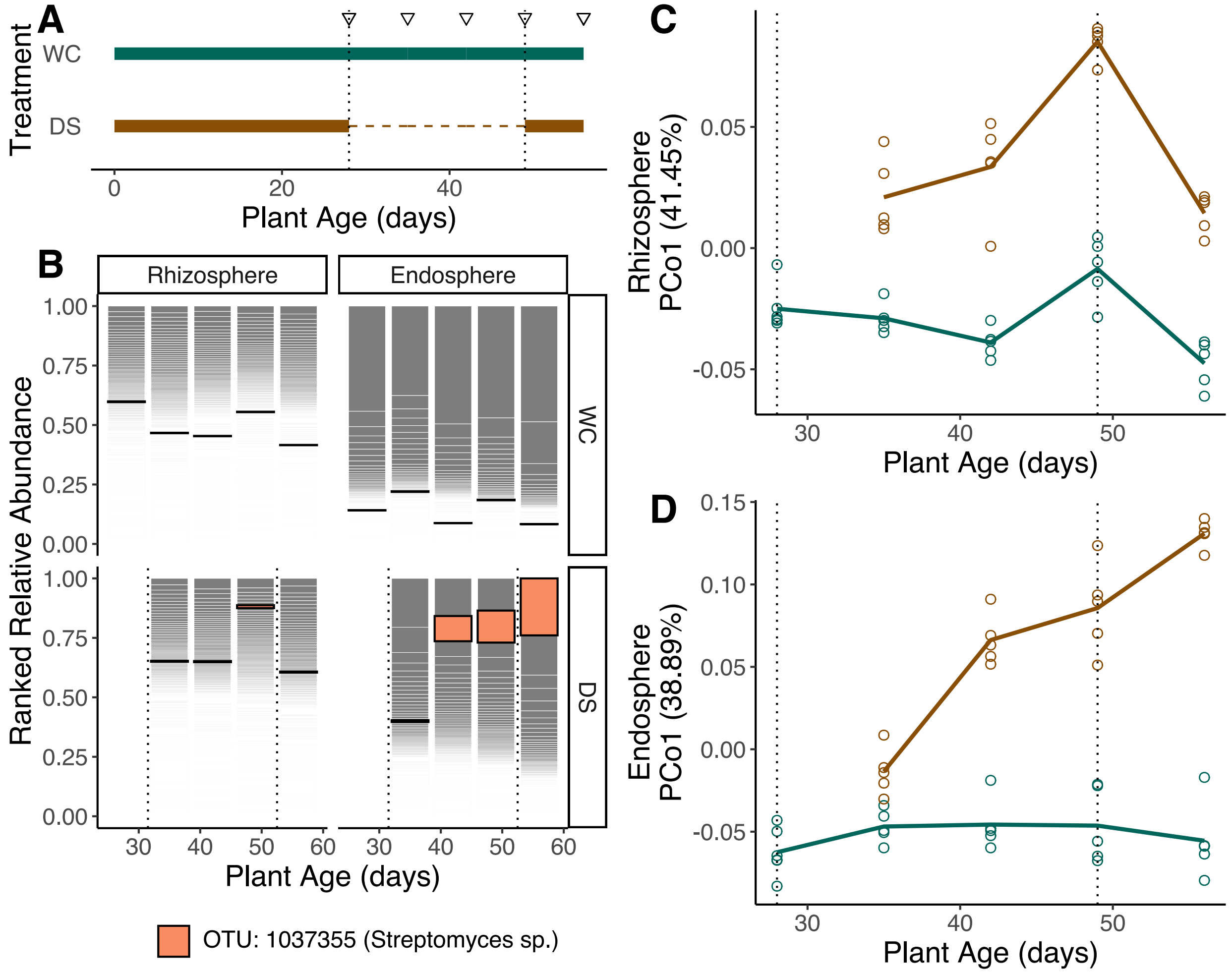


**Supplementary Figure 4.**

**OTU 1037355 displays reproducible trends in an independent experiment (A)** Timeline of the watering regimes followed by control (WC) and drought-treated (DS) plants. Horizontal lines represent the watering status during the experiment: solid segments indicate periods of constant irrigation while dotted segments indicate periods of suspended irrigation. Upside down triangles mark each of 5 collection time points. **(B)** Ranked relative abundances of individual community members throughout time. Each stacked bar plot displays all the OTUs detected in a particular time point: the height of individual bars represent the mean relative abundance of each OTU while the bar position across the y-axis indicates its rank within the community. The most abundant member of the semi-persistent enrichment module, *Streptomyces* sp. (OTU ID: 1037355), is highlighted. In all panels, the vertical dotted lines delimit the periods of suspended irrigation in each of the drought treatments. **C**, **D)** Beta-diversity patterns in the rhizosphere **(C)** and endosphere **(D)** communities. In both cases, the y-axis displays the position of each sample across the first principal coordinate (PCo) from a weighted UniFrac PCo analysis and the x-axis displays the age of the plant at the moment of sample collection. In all panels, the vertical dotted lines delimit the periods of suspended irrigation for the drought treatment.

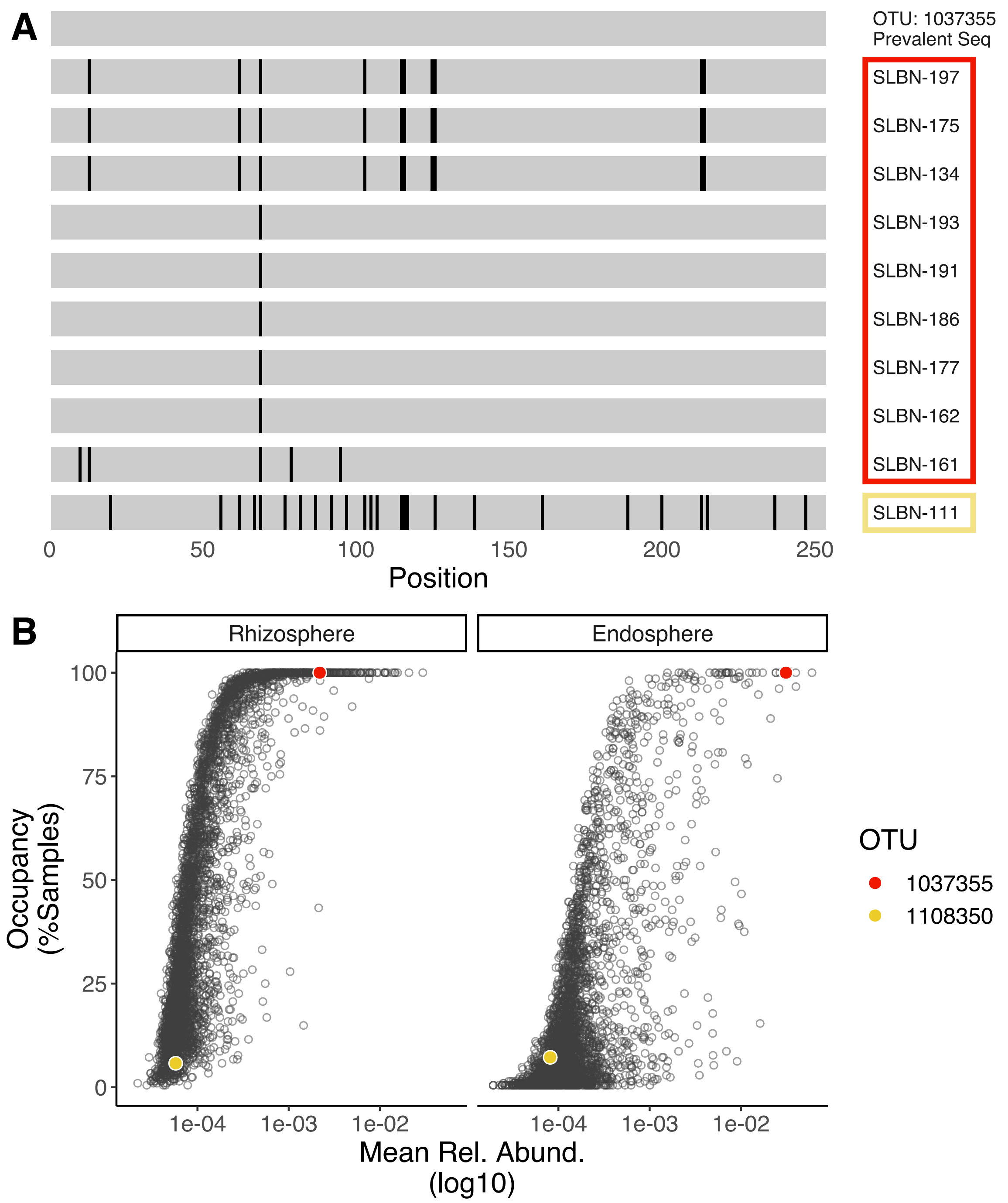


**Supplementary Figure 5.**
***Streptomyces* sp. SLBN-177 clusters with OTU 1037355**. **(A)** Sequence similarity along the V4 region of the 16S rRNA gene. The upper band represents the most abundant sequence of OTU 1037355 detected in the main drought experiment while the rest of the bands represent individual isolates collected from rice roots. Black lines indicate single nucleotide polymorphisms relative to OTU 1037355. All isolates highlighted in red cluster with OTU 1037355 at a 97% identity, while the isolate highlighted in yellow clusters with OTU 1108350. **(B)** Occupancy-abundance curves for the rhizosphere and endosphere communities. The x-axis displays the log-transformed mean relative abundance of each OTU while the y-axis displays the percent of samples in which each OTU was detected. OTU 1037355 and 1108350 are highlighted.

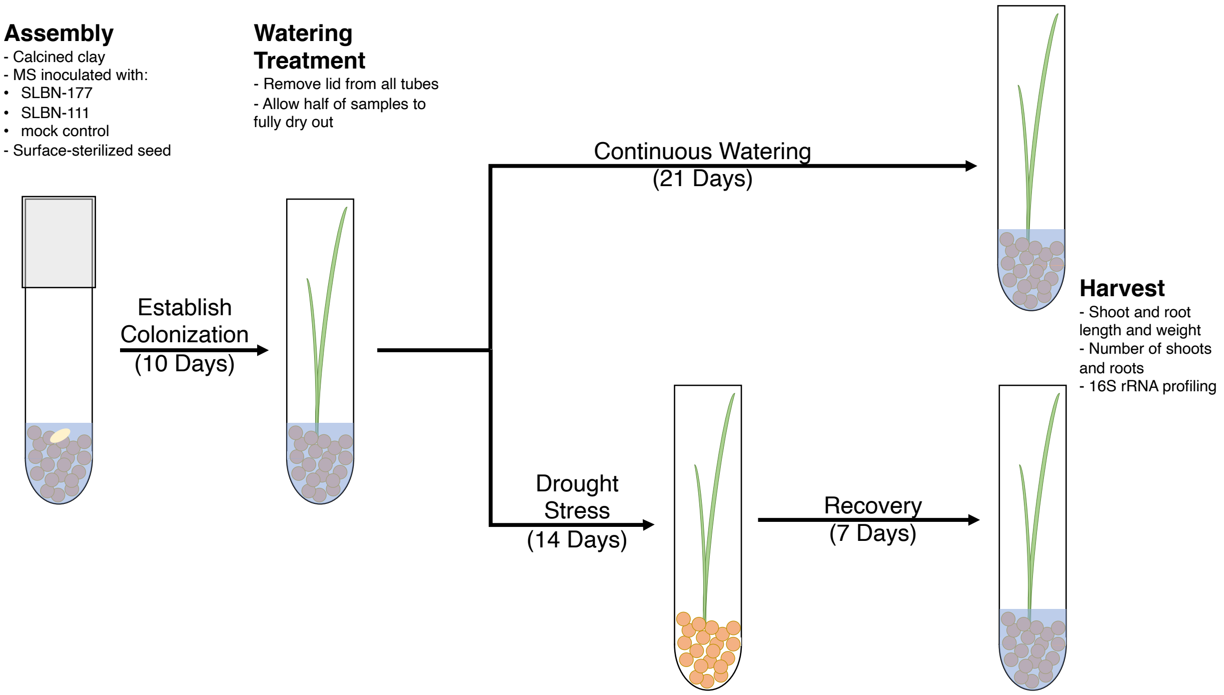


**Supplementary Figure 6.**

**Experimental setup for the phenotyping experiment.** Calcined clay, inoculum, and a surface sterilized seed were placed in 75 ml glass culture tubes, and were grown axenically for 10 days, followed by growth in an open system under drought or well watered conditions. This diagram corresponds to the timeline in figure 4A, and depicts an example tube under each condition at each stage of the experiment.

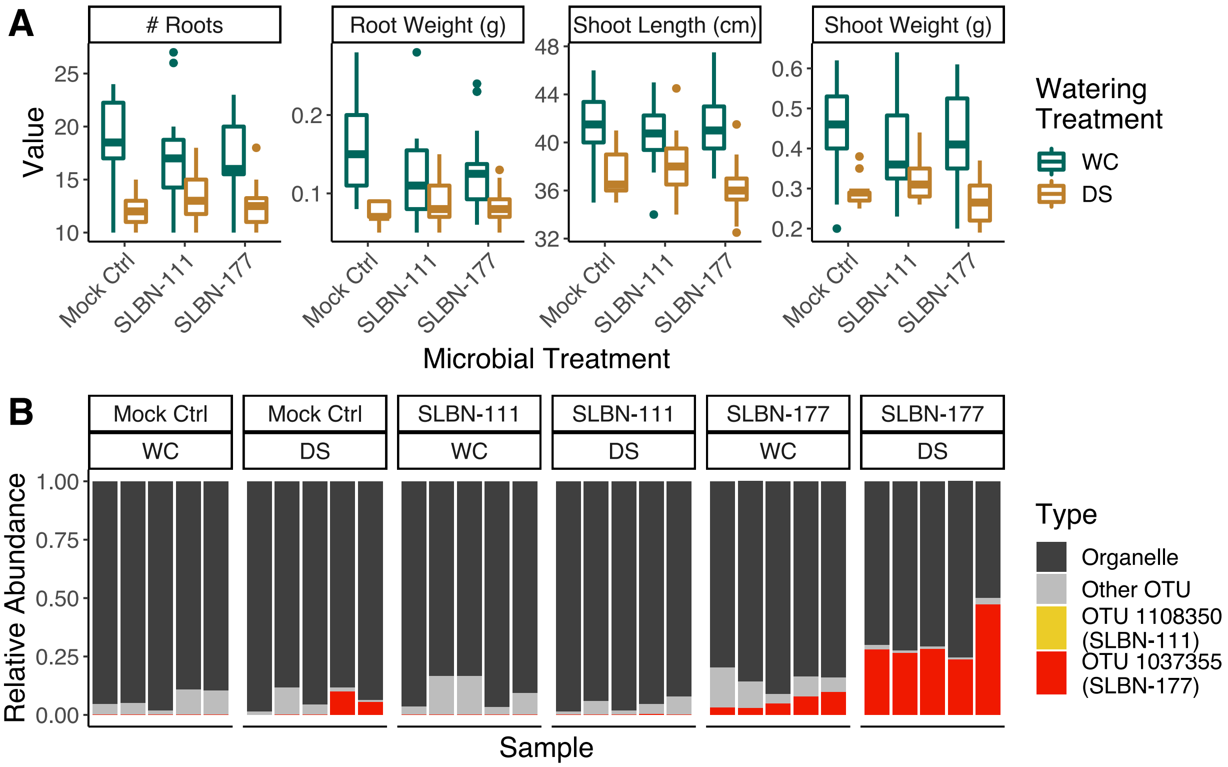


**Supplementary Figure 7.**

**Other phenotypic traits were not impacted by *Streptomyces* sp. SLBN-177. (A)** Distribution of measured plant phenotypes (number of roots, root weight, shoot weight, and shoot length) across microbial treatments and watering regimes. **(B)** Relative abundances of inoculated isolates in the endospheres of rice plants. Reads identified as mitochondria or chloroplast (collapsed as organellar reads and represented in black) were not discarded in order to measure the degree of colonization. Reads classified as OTUs other than 1037355 (SLBN-177) and 1108350 (SLBN-111) were collapsed and are represented in gray.

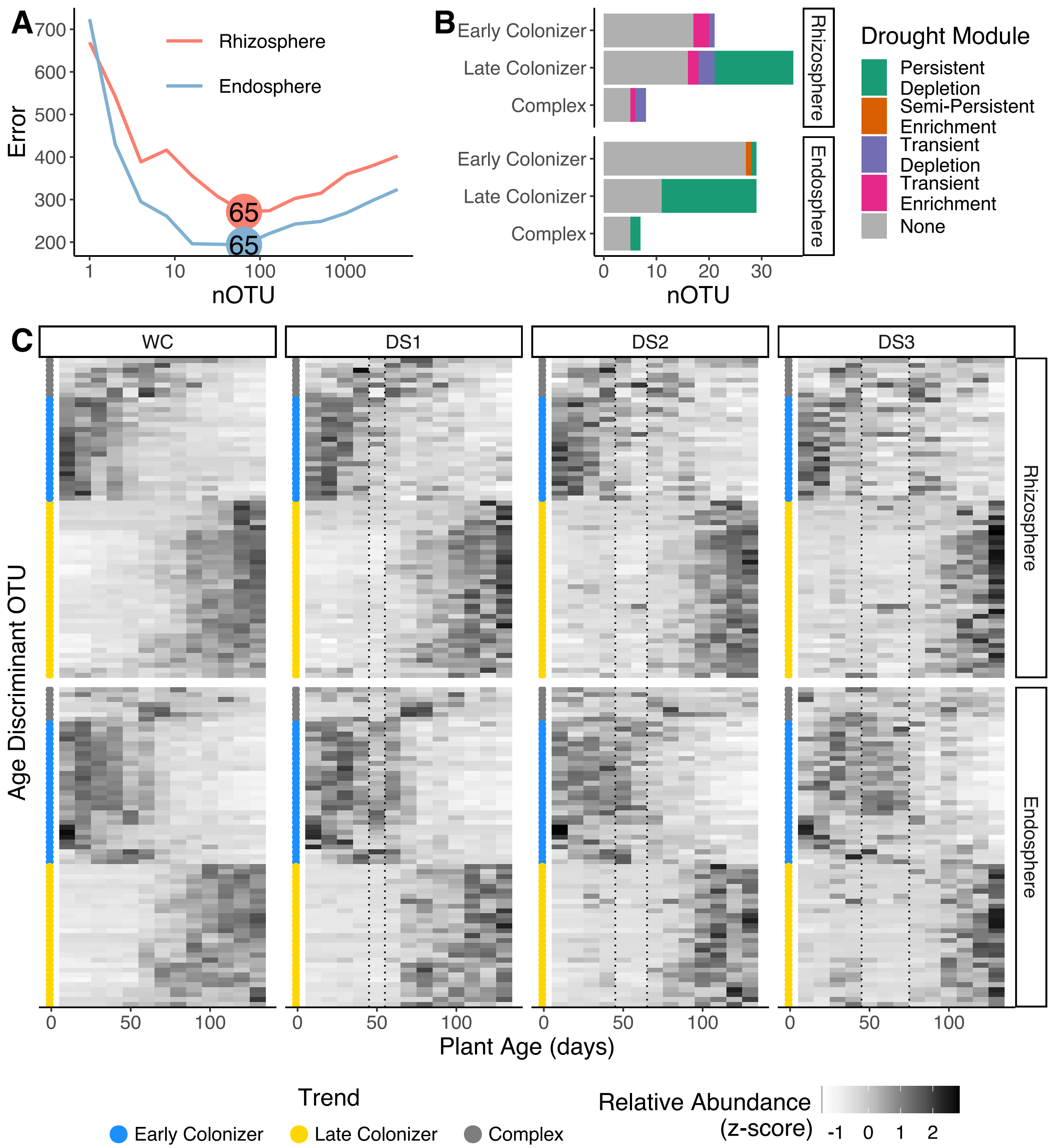


**Supplementary Figure 8.**

**Random forests can identify age discriminant OTUs. (A)** Cross validation error as a function of the number of OTUs used in each model. For both compartments, the lowest error was detected when using the 65 most important taxa. **(B) (B)** Overlap between the age-discriminant OTUs and the drought-responsive OTUs detected in each module (Figure 2B). **(C)** Hierarchical clustering of the relative abundances of the age-discriminant OTUs. The heatmap displays the z-transformed mean relative abundances of each OTU across drought treatments and time points. The colors at the left end of each vertical facet indicate the longitudinal trends exhibited by each taxa. In all panels, the vertical dotted lines delimit the periods of suspended irrigation for the drought treatment.

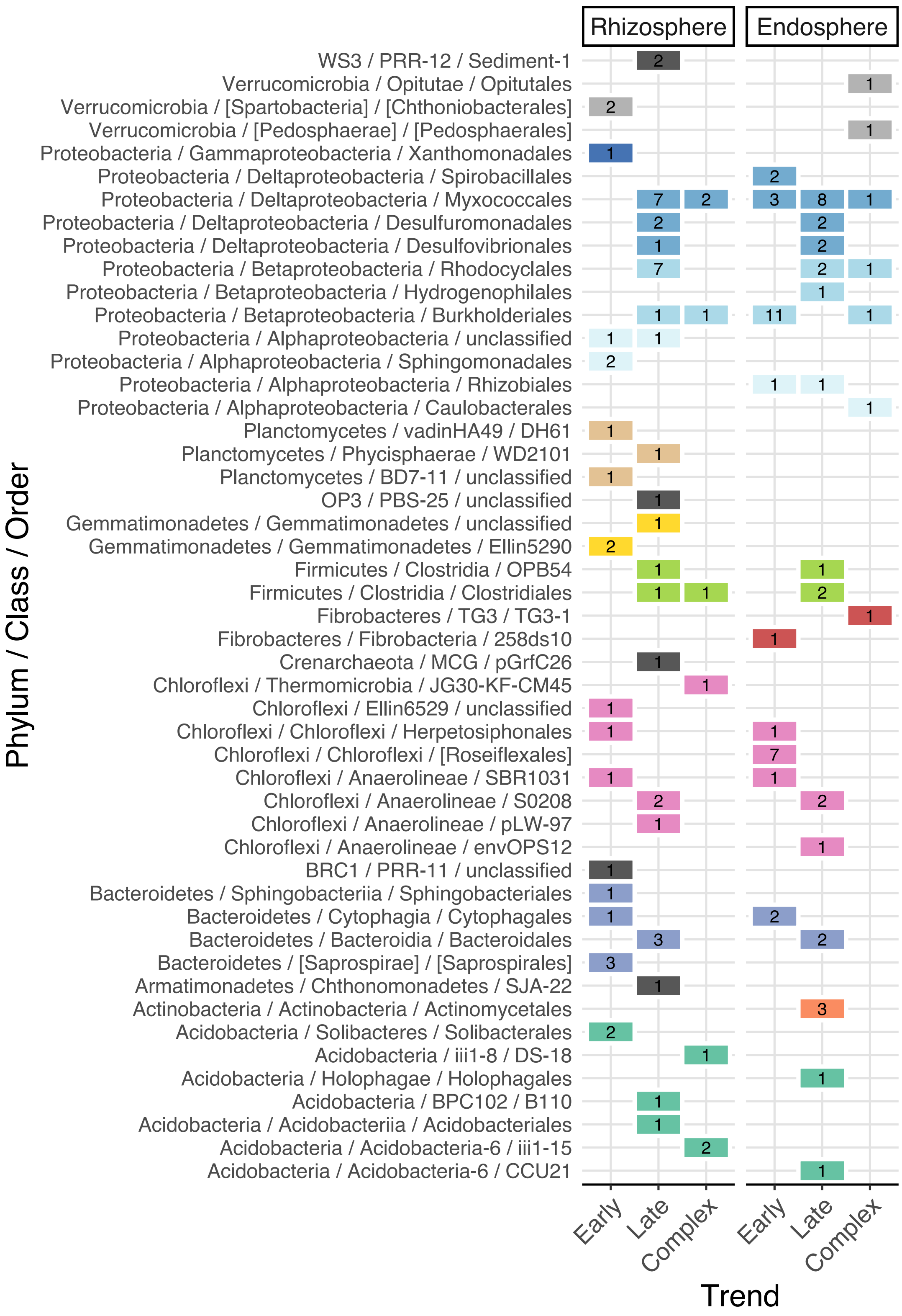


**Supplementary Figure 9.**
**Taxonomic classification of the age-discriminant OTUs in each of the longitudinal trends**. Each tile indicates the number of classified OTUs in a particular module, while the color represents membership to a specific Phylum / Proteobacteria class.

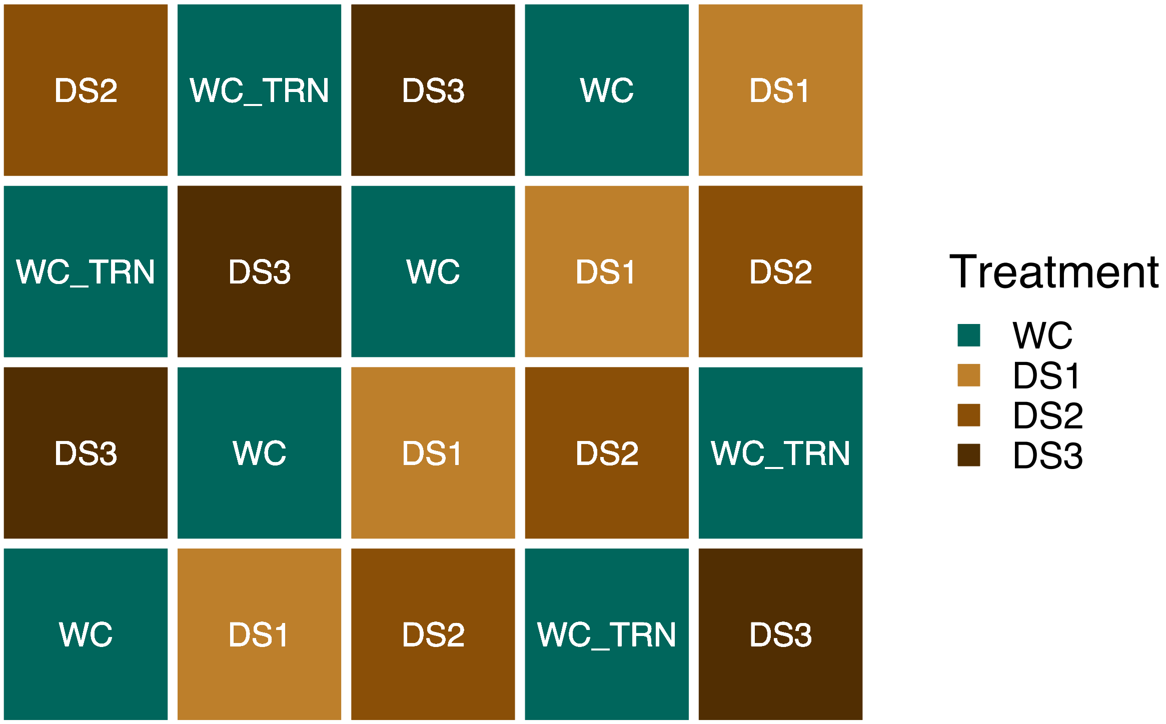


**Supplementary Figure 10.**
**Experimental design for the main drought experiment.** Randomized complete block design used for the greenhouse experiment. WC_TRN refers to the well-watered samples that were used to train the random forests models.

### Supplementary Tables

**Supplementary Table 1.**
OTUs with relative abundances significantly affected by drought treatments. Test statistics generated from Wald tests. Only significantly differentially abundant OTUs are present.

**Supplementary Table 2.**
Composition of clusters composed of differentially abundant OTUs as determined by Wald tests.

**Supplementary Table 3.**
ANOVA results on phenotypic traits measured in the inoculation experiment.

**Supplementary Table 4.**
Gene regions of potential secondary metabolite biosynthesis gene clusters as determined by antiSMASH. Only clusters with a significant BLAST hit to known clusters are present.

**Supplementary Table 5.**
Age-discriminant OTUs defined through random forest modelling, and their classification as late, early, or complex colonizers.
